# Advantageous prebiotic effects in ruminants of semi-refined chelates of dibasic cations with three- and four-carbon organic acids

**DOI:** 10.1101/2025.03.12.642899

**Authors:** Graeme D. Coles, R. John Pearce, Jacqueline S. Rowarth

## Abstract

Providing a semi-refined chelate of magnesium with mixed organic acids gave controlled magnesium supplementation of equivalent effectiveness to conventional uncontrolled administration of MgO in regard to prevention of milk fever in dairy cows. The supplement was palatable and supplementation produced no adverse health effects as adjudged by veterinary supervision.

Monitoring productivity of cows provided with the supplement compared with controls indicated that the use of the supplement provided a paranutritional improvement in feed utilisation, resulting in a significant increase in milk solids yield of up to 23% through the period of supplementation, together with an improvement in body reserves sufficient to sustain a continuing milk solids yield advantage of up to 5% for two months after cessation of feeding.

Since this increase in productivity was achieved without an increase in feed intake, use of the experimental supplement is considered to have resulted in a prebiotic effect in ruminant fermentation and consequently a commensurate reduction in methane intensity of production.

**Important Note on history of project:** The investigations reported in this paper were conducted under strict, independent veterinary supervision. This was because the responsible animal ethics committee first stated that the proposed study of the efficacy of a novel means of meeting magnesium supplemental requirements was of no academic interest, and refused to assess an application for ethics approval on those grounds. When the first experiment reported suggested the possibility of a prebiotic effect, the committee said that because there was no prior publication of such an effect in ruminants, the experimental hypothesis could not be sustained. This was despite the fact that prebiotic effects were well-known in humans and monogastric animals. As a consequence, the Committee refused to assess the research.

Subsequent investigations (e.g https://peerj.com/articles/18103) clearly indicate that the material under investigation could exert the prebiotic effect seen in monogastrics. In light of the present interest in inhibition of methanogenesis, the results reported below are of particular interest, requiring that they be placed in the public domain. Note that across both experiments, no adverse events related to administration of the prebiotic were observed by the veterinarian.

Conduct of the investigations reported has been compared with the ARRIVE guidelines, and found to be compliant.

## Introduction

Under commercial practice, magnesium supplementation is required when dairy cattle are managed on grazed pasture, particularly around parturition (Sansom et al. 1983). The most usual practice is to provide ∼20g/cow/day of elemental magnesium as MgO, either blended with a significant quantity of supplemental feed, or dusted on pasture. This is inefficient, since MgO is a poorly-available source of Mg, releasing between 14 and 20% of Mg supplied (Davenport et al. 1990), and is markedly unpalatable. The observed palatability of a semi-refined magnesium chelate in CDS (trademarked “Knewe®-Mg”) suggested that it might be a useful alternative to the normal commercial practice.

Fermentation of carbohydrate-rich plant materials with *Saccharomyces cerevisiae* is almost invariably intended to produce ethanol, but a number of co-products are also formed, including carbon dioxide, glycerol, lactic acid and other organic acids. These materials are products of the Embden-Meyerhof-Parnas metabolic pathway, used by *S. cerevisiae* to generate ATP, NADP and pyruvic acid, and these products are produced anaerobically (Waites et al. 2001). Production of glycerol and lactic acid are inevitable concomitants of ethanol production, as glycerol is synthesised by *S. cerevisiae* as a means of protecting the organism from the osmotic effects of ethanol, and lactic acid is synthesised from pyruvic acid when carbon dioxide content of the fermentation medium rises.

After fermentation is effectively complete, ethanol is distilled off, leaving *stillage* as a byproduct. Stillage consists of all the non-fermentable components of the original feedstock, plus the yeast biomass generated during fermentation, and the water and minerals used to suspend the original fermentation substrate(s). These components have considerable nutritive value for livestock (Iwanchysko et al. 1999; Mustafa et al. 2000), but are dilute, and are particularly attractive substrates for other microbial fermentations leading (unless intervention occurs), to rapid spoilage. It is commonplace to separate stillage into heavy and light fractions using centrifugation. The light fraction (“thin stillage”) consists largely of water, being between 2 and 4% solids, but contains salts and has significant BOD. It is therefore concentrated to “Distillers’ Condensed Solubles” (DCS), then co-dried with the heavy fraction (known as “wetcake”) to produce a material known as “Distillers’ Dried Grains and Solubles” (DDGS). This material is widely traded as a protein and energy source for ruminants, but its nutritive value is compromised by the heat required to convert the contained lactic acid and glycerol, which are naturally liquids (Thurmond and Edgar 1924; Borsook et al. 1933), into forms that can be dried.

Means of drying DCS with reduced impact on nutritive value have long been sought in commercial practice. Two of the present authors have developed a procedure for this (Coles and Pearce 2011), which involves converting the content of organic acids in thin stillage (and the liquid wetting the wetcake) to sparingly soluble chelates with dibasic cations. This provides a significant increase in the proportion of the content of the distillers’ grains and solubles that are dryable, and a similarly significant reduction in the level of components that require dehydration. The consequence is a reduction of at least 20°C in drying temperature, a significant reduction in drying time, and approximately 2/3 reduction in the amount of substrate required on which to dry the reacted CDS in conventional driers. It is also noted that provision of dibasic cations leads to the formation of crosslinked flocs of acidic, soluble protein, and these physical structures are thought to protect protein tertiary structure during heat processing, with the proteins being released in functional form from the flocs in the low-pH conditions of the abomasum when fed to ruminants.

Once dried, these semi-refined chelates are palatable to livestock and companion animals (Coles & Pearce, *op*.*cit*.), but it was not clear whether they would be useful sources of nutritionally important cations. Therefore, a case-control trial was conducted to see whether or not adequate magnesium supplementation could be achieved in commercial settings by feeding such DDGS treated and dried with a magnesium-rich source material.

It was hypothesised that sufficient Knewe®-Mg to supply 10g of elemental Mg daily would meet the needs of the periparturient dairy cow, and that this could be confirmed through lower incidence of magnesium deficiency symptoms compared to a control group conventionally supplemented. Such a study required also that other potential consequences in the health and productivity of the cows so supplemented in their feed be investigated.

## Materials and methods

A case:control investigation was carried out in animals intended to be milked commercially over winter. Following some unexpected observations, a second case:control investigation was carried out in a spring-calving herd, with feeding commencing about two months after mean calving date. A case:control design was implemented for the first trial, as animals could not act as their own controls in a crossover design, given the marked physiological changes occurring at calving. This approach was also used in the second trial.

### Animal ethics

In light of the GRAS status of the dietary interventions and the decision not to undertake invasive sampling procedures, the Animal Ethics Committee approached for advice declined to assess the proposed trial work for ethics approval. However, to satisfy the *amour propre* of the experimental team, independent veterinary supervision by Dr Michael Sheppard (Vet4Farm, Invercargill, NZ) was commissioned. This gave Dr Sheppard and his staff absolute authority to withdraw animals from the investigation, or to halt feeding if, in their view, the feeding was causing adverse effects. Throughout the two investigations described, this authority was not called upon.

The conduct of the experimental procedures reported here has been compared with the ARRIVE guidelines (Percie et al. 2020), and found to be compliant.

### Investigation 1

#### Experimental product

Unless otherwise stated, all materials included in animal diets were of feed grade.

Experimental product was prepared according to Coles and Pearce (op. cit.), modified as follows.

Distillers Condensed Solubles (DCS) syrup was collected from normal production from the facilities of Shoalhaven Starches Pty Ltd at Bomaderry, New South Wales, Australia, and transferred to the premises of Halcyon Products Pty Ltd in Melbourne, Victoria, Australia. Under normal conditions, DCS from this source contains approximately 25% lactic acid (dmb), but this shipment was somewhat depleted in this material, so was supplemented with technical grade lactic acid (All Raw Materials Pty Ltd, Young, NSW, Australia).

DCS was heated to 55°C, and reacted with commercial feed grade magnesium oxide (Causmag International Pty Ltd, Melbourne, Victoria, Australia) with continuous stirring and monitoring of pH. When pH reached 7, addition of magnesium oxide ceased, and the product was prepared for drying. Small-scale spray drying studies had revealed the need to incorporate a quantity of maltodextrin to improve flow properties, so 10% additional maltodextrin was mixed with the slaked DCS.

The resulting mixture was spray dried using conventional techniques, and packaged for storage and transport in 20 kg quantities in moisture-proof plastic bags in cartons. Its composition is given in Table 1.

**Table 1.** Composition of prepared Distillers’ Condensed Solubles (Dairy One, Inc, Ithaca, New York, USA).

| Component | Content as fed |
| --- | --- |
| Moisture (%) | 10.5 |
| Drymatter (%) | 89.5 |
| Crude protein (%) | 16.7 |
| Available protein (%) | 13.6 |
| ADICP (%) | 3.0 |
| Adjusted crude protein (%) | 14.5 |
| Acid detergent fibre (%) | 1.8 |
| Neutral detergent fibre (%) | 4.2 |
| Nonfibre carbohydrate (%) | 58.2 |
| Starch (%) | 0.5 |
| Water-soluble carbohydrates (%) | 20.1 |
| Crude fat (%) | 1.9 |
| Ash (%) | 9.44 |
| Total digestible nutrients (%) | 68.0 |
| NE<sub>L</sub> (Mcal/kg) | 1.57 |
| NE<sub>M</sub> (Mcal/kg) | 1.62 |
| NE<sub>G</sub> (Mcal/kg) | 1.06 |
| Calcium (%) | 0.11 |
| Phosphorus (%) | 0.55 |
| Magnesium (%) | 2.82 |
| Potassium (%) | 0.93 |
| Sodium (%) | 0.782 |
| Iron (ppm) | 115.0 |
| Zinc (ppm) | 26.0 |
| Copper (ppm) | 6.0 |
| Manganese (ppm) | 57.0 |
| Molybdenum (ppm) | 1.3 |
| Sulphur (%) | 0.21 |
| Chloride ion (%) | 0.59 |
| pH | 8.6 |
| Dietary cation-anion difference (mEq/100g) | 31.0 |

Apparent digestible energy was estimated by NIR to be 14.4 MJ/kg.

#### Feed supplement manufacture

##### Experimental diet

For the purposes of this experiment, it was estimated that magnesium availability from the experimental supplement would be similar to availability of magnesium from magnesium chloride or sulphate, i.e. approximately 65%. Provision of 6.5 g of magnesium absorbed therefore required daily intake of sufficient experimental supplement to provide a total of 10 g of magnesium, and this quantity was available from 400 g of the spray dried product.

For convenience of feeding, the 400 g of experimental supplement was incorporated into 2 kg of diet, the balance consisting of 1600 g of ground New Zealand-grown new season’s winter wheat (12.2% protein dmb). This mixture was supplemented by the manufacturer (Seales Winslow Ltd, Tinwald, New Zealand) with 0.1% of the manufacturer’s proprietary palatability enhancer. 19.9 t of this diet was manufactured and fed.

##### Control diet

The control diet consisted of 40 g of dairy nutritional grade magnesium oxide (Causmag International Pty Ltd, Melbourne, Victoria, Australia) in 2 kg of supplement. The supplement consisted of the same wheat as used in the experimental diet, augmented by sufficient soybean meal (46% protein dmb) (Viterra Ltd, Auckland, New Zealand) to compensate for the difference in the protein content between the wheat and the experimental supplement displaced from the experimental diet formulation. This diet was also supplemented with the same proprietary palatability enhancer. A total of 119.0 t of this diet was manufactured and fed.

#### Composition

Composition of the two supplements was determined by an independent laboratory (Nutrition Laboratory, Institute of Food, Nutrition and Human Health, Massey University, Palmerston North, New Zealand). Results are given in Table 2.

**Table 2.**
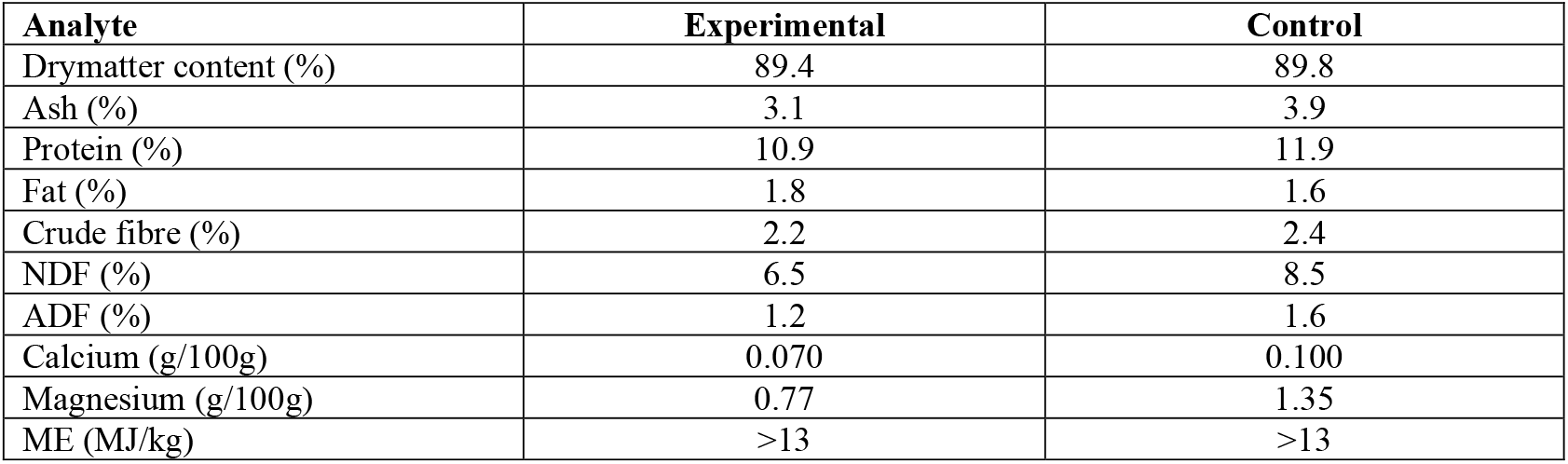
Supplement composition.

#### Animals

The trial involved a winter milking herd of New Zealand Friesians owned by Southern Centre Dairies Ltd. The dairy platform is located at East Limehills, just north of Winton in Southland, New Zealand. Approximately 800 cows, calving from February to April, were milked twice daily. Prior to the commencement of the trial, dry cows were conventionally grazed on pasture with supplementation with conserved forage when necessary, and magnesium supplementation was supplied in the form of magnesium oxide broadcast onto pasture.

Within this herd, a mating block of 180 third and fourth parity cows, due to start calving on March 18^th^ 2012, was selected for detailed investigation. On March 14th, these cows were separated into two groups as follows. All the cows were brought into the yard of the milking platform, and allowed to make their normal social arrangements. Cows were then run on to the rotary platform in the order that suited them. This order was recorded, and cows allocated to either the control group or experimental group alternately (the “condition” groups). After the 30th cow had come off the platform each fifth cow was alternately allocated to further smaller experimental groups (sampler cows). In this way, two groups of 80 condition cows and two groups of 10 sampler cows were created. The process was intended to provide a random allocation of cows to groups by splitting an entire population in half, and abstracting a further subpopulation at random.

As the animals came off the rotary platform, their ear tag numbers were recorded in order. Sampler cows were provided with coloured ear tags (blue for experimental diet, red for control diet), and drafted out of the main block. The condition cows were then individually judged for body condition by a veterinarian (Anon. 2012), and returned to grazing. Cows whose body condition was below 4 were excluded from the trial. The sampler cows were allowed a period on pasture, and then returned to the platform for sample collection prior to going back to the main mob.

Allocation of animals to groups was recorded in the herd management database, and these allocations were used thereafter to determine which of the two dietary supplements each animal was fed, and to aid drafting at each sampling visit.

#### Measurements

Immediately after each cow calved and entered the milking herd, milk yield was recorded daily. For various reasons, approximately 30 animals from the mating block of 180 cows did not enter the milking herd during the 3 weeks expected. This was largely due to cows not conceiving in the first round of mating. The final numbers of cows, by treatment, were 65 control cows, 66 cows consuming the experimental diet with 9 cows in each of the subsample groups.

Once all cows had entered the milking herd, the subsample animal were retained after milking each 14 days, and returned to the milking platform to provide a milk sample for analysis. Milk samples were frozen and shipped to a commercial milk testing laboratory.

#### Data processing

As milk samples were collected, they were forwarded to an independent laboratory (MilkTest Ltd, Hamilton, New Zealand) for analysis. Analytes were:

- Milk fat concentration;
- Milk protein concentration;
- Milk urea concentration, and;
- Somatic cell count.

From these individual analytes, it was possible to estimate total milk solids concentration and yield, and total milk N concentration and content. It is known that milk lactose yield is very highly correlated with milk volume, because the osmotic effect of the lactose synthesised in the udder is to draw water into the lumen of the udder alveolus (Haile-Mariam and Pryce 2017). Therefore, using a standard set of values for milk solids composition (USDA food composition database) and the daily milk volume obtained for each cow, her lactose secretion could be estimated.

On completion of the feeding period, daily herd milk records were downloaded from the herd data management system and sorted according to treatment in an Excel spreadsheet. Data were then transferred to MiniTab v14 for analysis of variance using a Generalised Linear Model (GLM-ANOVA). Such a model allows for lack of balance in the data, particularly at the beginning of the experimental period, when the number of cows being milked each day grew as calving progressed.

#### Empirical results

The number of animals suffering milk fever, the number of animals with slow onset of milk production, and the number requiring treatment for disorders such as mastitis were reported by the herd manager, but were so low as not to be amenable to statistical analysis.

### Investigation 2

In this investigation, preliminary indications of a paranutritional (prebiotic) effect of the experimental product were further evaluated.

#### Experimental product

Spray drying proved too costly to be commercially feasible, so the experimental product for this trial was manufactured and blended with millrun (a mixture of wheat bran and pollard in the proportions normally produced during flour milling). Experimental investigations revealed that a mixture consisting of equal proportions of millrun solids and DCS solids reacted with MgO could be efficiently dried in a commercial drum drier, so a blend of this composition was manufactured for trial purposes. The outcome of the first investigation was that supplying Knewe®-Mg at a rate providing 8g/day of Mg as lactic acid chelate was sufficient to meet the dairy cow’s Mg requirements, with virtually no incidence of milk fever^1^. Therefore, the Knewe®-Mg, co-dried with mill run, was able to meet Mg supplementation requirements when fed at a rate of 800g/day. For the second investigation, this quantity of Knewe®-Mg was blended with small amounts of additional minerals and a palatability enhancer. A control supplement was manufactured, based on wheat wholemeal and MgO, together with the same blend of additional supplementary minerals. Note that the palatability enhancer was required to mask the bitterness of the MgO in the control diet, so was not needed in the experimental treatment. However, to ensure proper control, it was included in both treatment arms.

#### Experimental design

The trial herd used included 450 multiparous animals. These animals were sorted according to their herd accession number, then allocated at random to three equal-sized treatment groups. Based on the results of trial one, a power calculation showed that subsampling groups of seven animals were sufficient, selected at random from each treatment group.

Three experimental diets were used: the control product, the experimental product and a 50:50 blend of the control product and the experimental product (“Blend”). Animals were provided with 500g of the diet for the trial arm in which they were included at each of two milkings daily, providing a total of 1000g of supplement daily. Animals in the herd not included in the trial groups were fed 1000g of the control supplement daily. Feed acceptability was high, with >99% complete consumption during the ∼ 10-minute milking period.

The basal diet for all animals was grazed ryegrass:white clover pasture, supplemented during a period of (relative) drought with ryegrass:white clover silage conserved from the previous autumn. This silage was fed under cover through a “long wire” allowing all animals equal access to feed. Feed allocation was carefully monitored to avoid wastage, but there was no indication of variation in animal appetite between treatments.

#### Sample collection

Fortnightly milk sample collections were made from the subsampling groups. These groups were drafted from the main herd after the evening milking from the previous day, and grazed overnight on a separated pasture allocation next to the main herd. The subsampling treatments were brought to the milking facility after the rest of the herd, brought on to the rotary platform together, and provided with their supplement allocation. Each animal was milked separately into a single 20L bucket. Milk was thoroughly mixed and subsampled and labelled for analysis. Milk produced in excess of the bucket capacity was captured by the milking shed system, but the main sample was discarded to waste. Milk quality parameter values were determined by an independent laboratory (MilkTest Ltd, Hamilton, New Zealand).

#### Data processing

Data were processed as for trial 1. Daily milk yield (three-day moving average) was determined for each treatment and used to calculate fortnightly values for daily production of milk fat, milk protein and milk urea. Fortnightly estimates of somatic cell count were log-transformed before analysis. This was because on a whole-herd basis, minor changes are due to major rises in level in one or a few animals, so if such a rise happened to occur in a single subsampling animal, eliminating it from analysis would undermine the power of the experiment.

## Results and discussion

### Experiment 1

#### Anecdotal observations

The primary goal of this experiment was to test the palatability and effectiveness of Knewe®-Mg as a magnesium supplement. Anecdotal evidence (T. Mead, herd manager, *pers. comm. 2012*) was that almost no evidence of differential incidence of milk fever (periparturient paresis (Barrington 2011)) was observed. A total of four animals in each treatment arm showed transient symptoms, insufficient to require intervention, while a single animal in the control arm exhibited symptoms during a presentation requiring veterinary attention. Therefore, this trial was unsuccessful in showing a differential **benefit** of Knewe®-Mg for milk fever management in what was apparently a very well-managed herd.

Other empirical observations included that animals fed the experimental product tended to come into milk more rapidly than control-fed animals, a finding of significance for neonatal health.

#### Subclinical effects

The purpose of this study was to assess the relative value of Knewe®-Mg and a control treatment for providing effective magnesium supplementation. The primary endpoint showed that Knewe®-Mg was as effective as the conventional management approach, but analysis of productivity was undertaken to determine whether the product had any subclinical effect. Figure 1 shows the relative productivity of control-and experimentally fed animals over the first 3 months of lactation.

**Figure 1.**
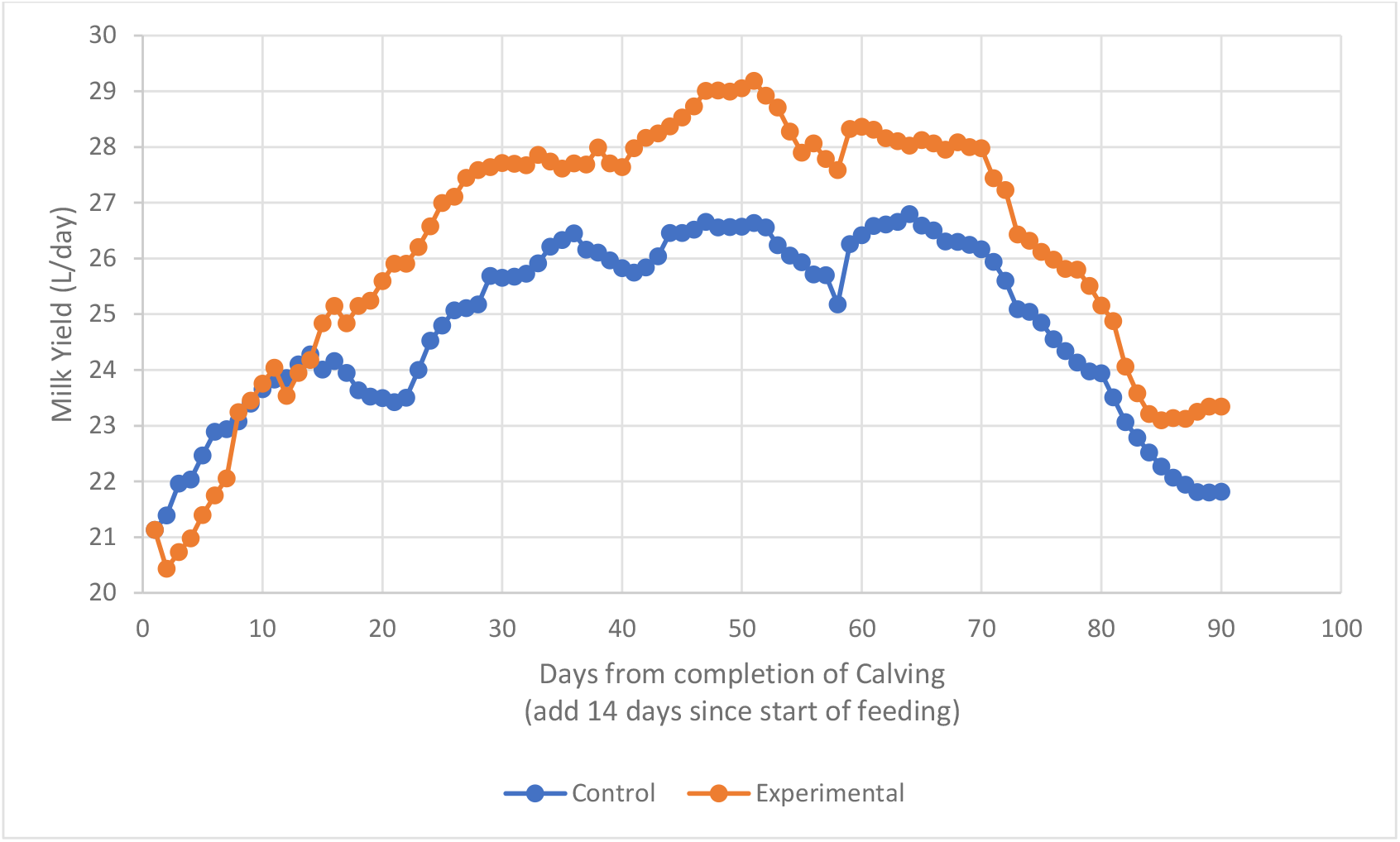
Daily milk production per day per cow for the first three months of lactation in trial one.

Experimentally fed animals were initially less productive than the control-fed animals (Figure 1), but soon surpassed the control daily milk yield. Across the entire trial period, the uncorrected yield advantage of the experimentally fed animals was 6%. When milk composition was also considered, a further productivity advantage was observed. Milk protein was slightly elevated, and there was a marked increase in milk fat content. However, collection of samples less than 90 minutes following complete milking led to concentration values above the range normally found in *Bos taurus*, so only relative values were used to estimate overall advantage in productivity. This estimate, of 16%, combining milk yield and milk solids advantages, was sufficient to stimulate interest in a second experiment in which milk composition and component yield were the primary endpoints of interest.

### Experiment 2

This experiment was conducted in a spring-calving herd, and used a larger number of animals, in three treatments.

### Baseline analyses

To ensure that the trial groups and subgroups were representative samples of the herd as a whole, data collected at randomisation for the parameters measured were subjected to separate statistical analysis.

#### Milk yield and composition

Data for the first three days of the trial were analysed.

#### Milk Composition

Data obtained from the first sample collection were analysed using one-way ANOVA. Results are given in Table 3. Note that NZ solids is the sum of Fat % and Protein %. Milk energy is calculated as the sum of lactose% + 2.25 x Fat%. (The dietary energy content of fats is 2.25 times greater than carbohydrate on a weight for weight basis.) Nitrogen (N) ratio is the product of urea concentration divided by protein concentration, with lower values considered desirable.

**Table 3.** Baseline performance measurements.

| .Character | Experimental diet<br>(% Control) | Blend<br>(% control) | Control diet | Significance (p) |
| --- | --- | --- | --- | --- |
| Milk yield (L/day) | 29.13 (100.9) | 28.27 (97.9) | 28.87 | 0.132 |
| *Milk Composition* |  |  |  |  |
| Fat (%) | 6.73 (120.8) | 6.23 (111.7) | 5.57 | 0.606 |
| Protein (%) | 3.60 (108.1) | 3.40 (102.1) | 3.33 | 0.518 |
| Lactose (%) | 4.58 (95.2) | 4.67 (97.1) | 4.81 | 0.572 |
| NZ Solids (%) | 10.76 (120.1) | 9.63 (108.1) | 8.91 | 0.301 |
| Urea (mM/L) | 17.76 (103.7) | 20.79 (121.4) | 17.12 | 0.607 |
| Log SCC* | 5.15 | 4.91 | 5.79 | 0.155 |
| Nitrogen (N) ratio | 5.06 (98.0) | 6.26 (122.0) | 5.13 | 0.582 |
| Milk Energy | 20.72 (119.4) | 18.68 (107.6) | 17.36 | 0.405 |

\*Ratio to control is not appropriate when reporting transformed data

Despite substantial absolute and relative difference for most characters between groups, the within-group variability is such (Gross 2022) that the group results are not significantly different for any character at baseline. Consequently, if differences in any character achieve statistical significance in the course of the trial, we assume adequate *prime facie* evidence for a real effect. No attempt was made, therefore, to use any baseline analyte value as a covariate in subsequent analyses.

#### Milk component yield

Milk component concentration variability may be due to variation in the absolute amount of components secreted, or the amount of diluent in which the components are dispersed. Therefore, **absolute** amounts of milk solids produced each day for each individual animal were calculated and subjected to ANOVA.

Table 4 shows that the mean group performance for the control and the experimental diet at baseline are essentially indistinguishable, and while the apparent performance of the “Blend” group is lower, the difference is not at all distinguishable statistically-speaking.

**Table 4.** Comparison of absolute milk solids yield at baseline.

| .Character | Experimental diet<br>(% Control) | Blend<br>(% control) | Control diet | Significance (p) |
| --- | --- | --- | --- | --- |
| Milk yield (L/day) | 29.13 (100.9) | 28.27 (97.9) | 28.87 | 0.132 |
| NZ Solids (%) | 2.801 (98.3) | 2.538 (89.1) | 2.849 | 0.856 |
| Mean days after calving | 34.0 | 42.78 | 44.125 | 0.014 |

Note that the mean days in milk for the animals in the trial group is significantly **lower** than the same parameter value for the other groups. This further supports the contention that the **performances** of the three experimental groups is not significantly different to each other at baseline. Because of missing values among the milk composition data, no rational set of covariates could be used.

### Trial analyses

#### Milk yield

For various management reasons, a small number of cows were removed from the herd during the trial period, leading to unbalanced data. Milk yield data from cows not completing the full feeding regimen were excluded from analysis. Daily yield data for individual cows were analysed using the GLM procedure in Minitab (version 16). Milk yield profiles for the three treatments are given in Figure 2. To exclude the possibility of a systematic bias in milk yield, the mean yield over the first three days of the trial for each cow was used as a covariate in the analysis. Consequently, the estimate of daily milk yield advantage is conservative.

**Figure 2.**
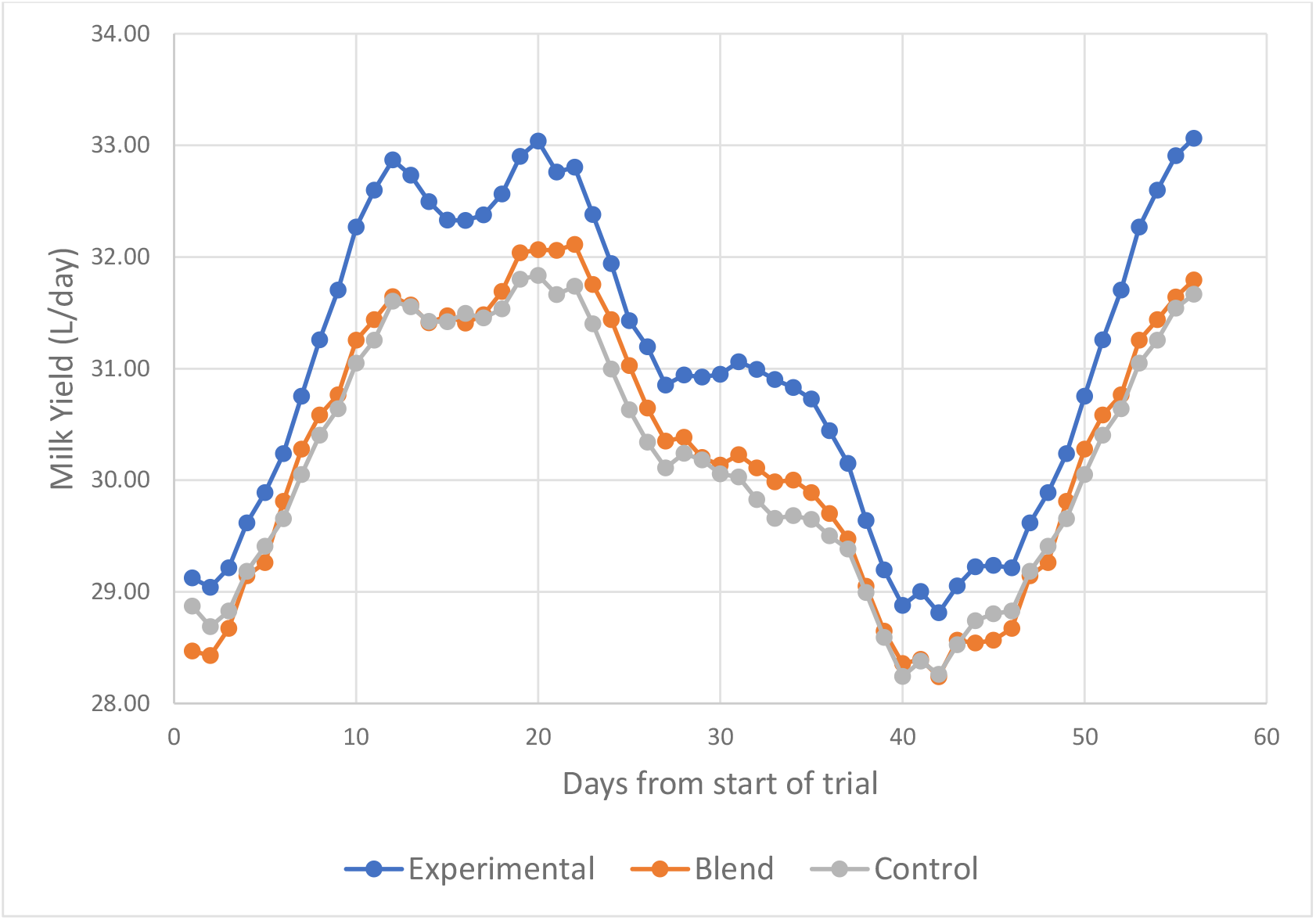
Comparison of daily milk yield for the three treatments included in the second trial. The decline in milk yield from early December to the beginning of January corresponded to an extended period of hot dry weather, during which the herd was supplemented with palm kernel expeller meal (PKE). A single 50mm rainfall event in early January was followed by application of urea and dramatic pasture recovery, leading to a return to expected milk yield.

Thus, using initial milk yield as a covariate, the cows fed the control diet produced mean 30.06 L/day, the animals fed the blend, 30.44L/day (101.3% of control) and the animals fed the experimental diet 30.75 L/day (102.3% of control) (p = 0.000).

Total milk production for the period of the trial was also calculated, and subjected to one-way analysis of variance. Results were: Control diet, 1635.5L; Blend, 1622.4L (99.2%); Experimental diet, 1700.6L (104.0%); p=0.169. As indicated by the previous study, the differences in the summed milk yields were slightly higher than the differences in mean daily yields, but the accumulation of data to produce the summed yield greatly reduced the number of degrees of freedom for the analysis, and concealed a small but statistically-important treatment:time interaction.

#### Milk Yield: sampler cows

The milk yield data analysed above were derived from the main trial groups, each consisting of approximate 140 cows. Data from groups managed separately for provision of milk for composition analyses were also analysed. A larger yield advantage was achieved (Control diet, 28.63L/day; Blend, 26.96L/day (94.2%); Experimental diet, 29.98L/Day (104.7%) (p=0.000)), and when mean milk production/day for the first three days was used as a covariate, the advantage to the experimental diet was increased (Control diet, 27.73L/day; Blend, 28.31L/day (102.1%); Experimental diet, 29.46L/day (106.2%) (p=0.000)).

#### Milk Composition

Milk composition data were analysed as for milk yield, using a factorial analysis in GLM in Minitab (Table 4) with individual animal responses nested within treatments. There was no significant interaction between treatment and sampling date for any character. Because of the known variability in milk component concentration immediately after calving (e.g. Gross (2022)), no analysis of covariance using baseline data was conducted.

As noted above (Tables 3 and 4), the baseline means for treatments, although variable, were not significantly different. The variability that developed during the 2^nd^ trial therefore demanded closer investigation.

#### Milk analyte yield

Using milk volume produced by each cow at each sampling time, milk analyte **yield** was obtained from milk analyte concentration (Table 6).

It was noted that somatic cell count varied significantly between treatments, so the log-transformed measure of somatic cell count was used as a covariate in a generalised linear model ANOVA for milk solids yield. Results were: Experimental, 2.393kg/day; Blend, 2.044kg/day; control, 2.159kg/day; p=0.014. These results indicate that somatic cell count, an indicator of mastitis incidence, affected milk solids yield, and if its effect is accounted for, milk solids yield rises by almost 7%.

### Changes in Milk Solids yield with time

Milk solids yield changes with time were determined from the sampler cows. Data providing the results in Table 4 were analysed, and results from each sampling time are shown in Figure 3.

**Figure 3.**
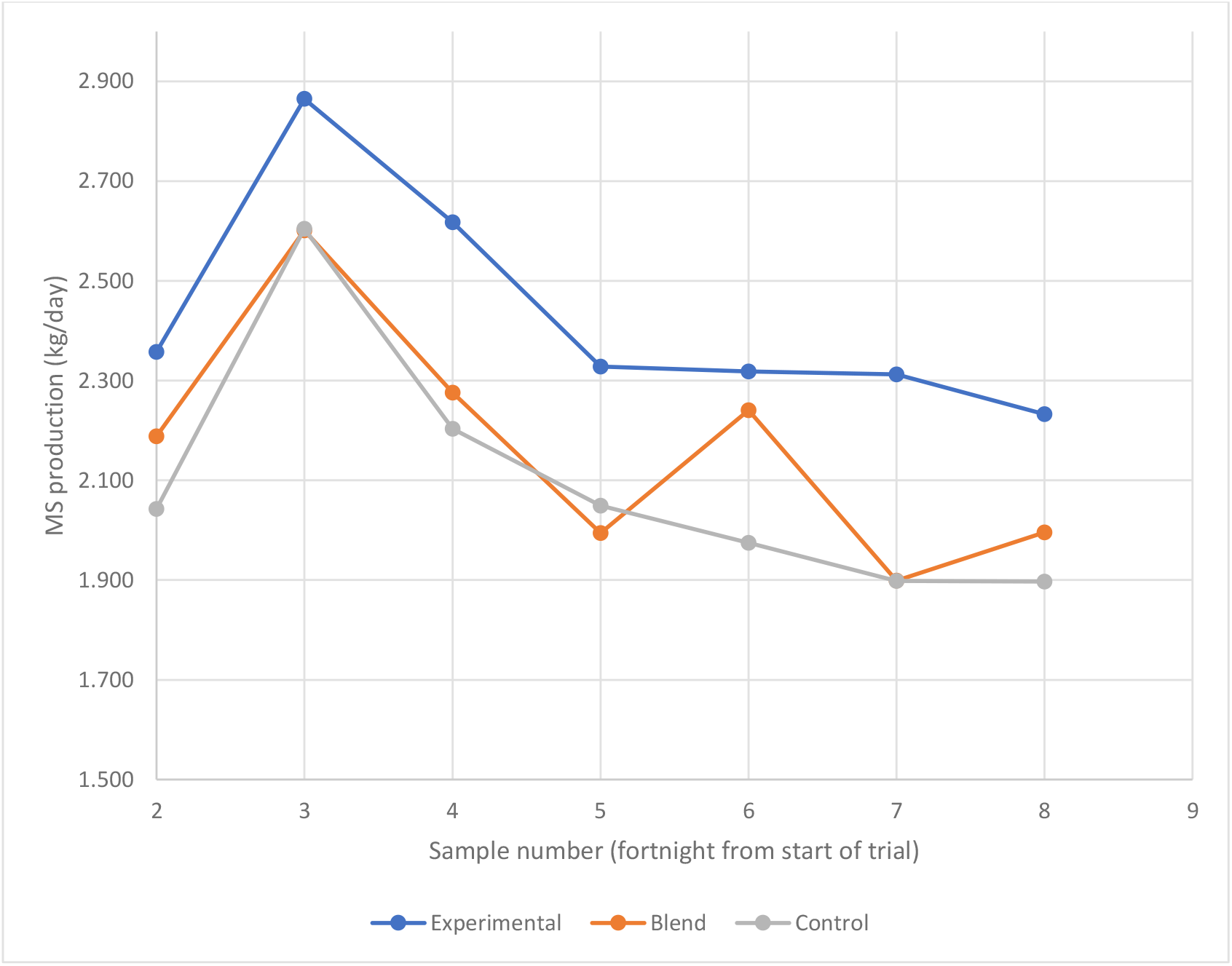
Milk Solids yield change with time by experimental treatment in Trial 2. Data plotted are GLM estimates of treatment x sampling date means.

The prebiotic effect was displayed only in animals being fed a full daily dose, as determined by the provision of an adequate amount of elemental magnesium. As noted above, it appears that when commencing feeding of the prebiotic supplement **after** calving, most, if not all, of the increase in milk solids yield was derived from additional milk fat secretion.

Therefore, an analysis of the relative production of milk protein and milk fat was conducted by comparing the ratios of the concentrations of milk fat to milk protein (F/P ratio) throughout the trial. (Figure 4).

**Figure 4.**
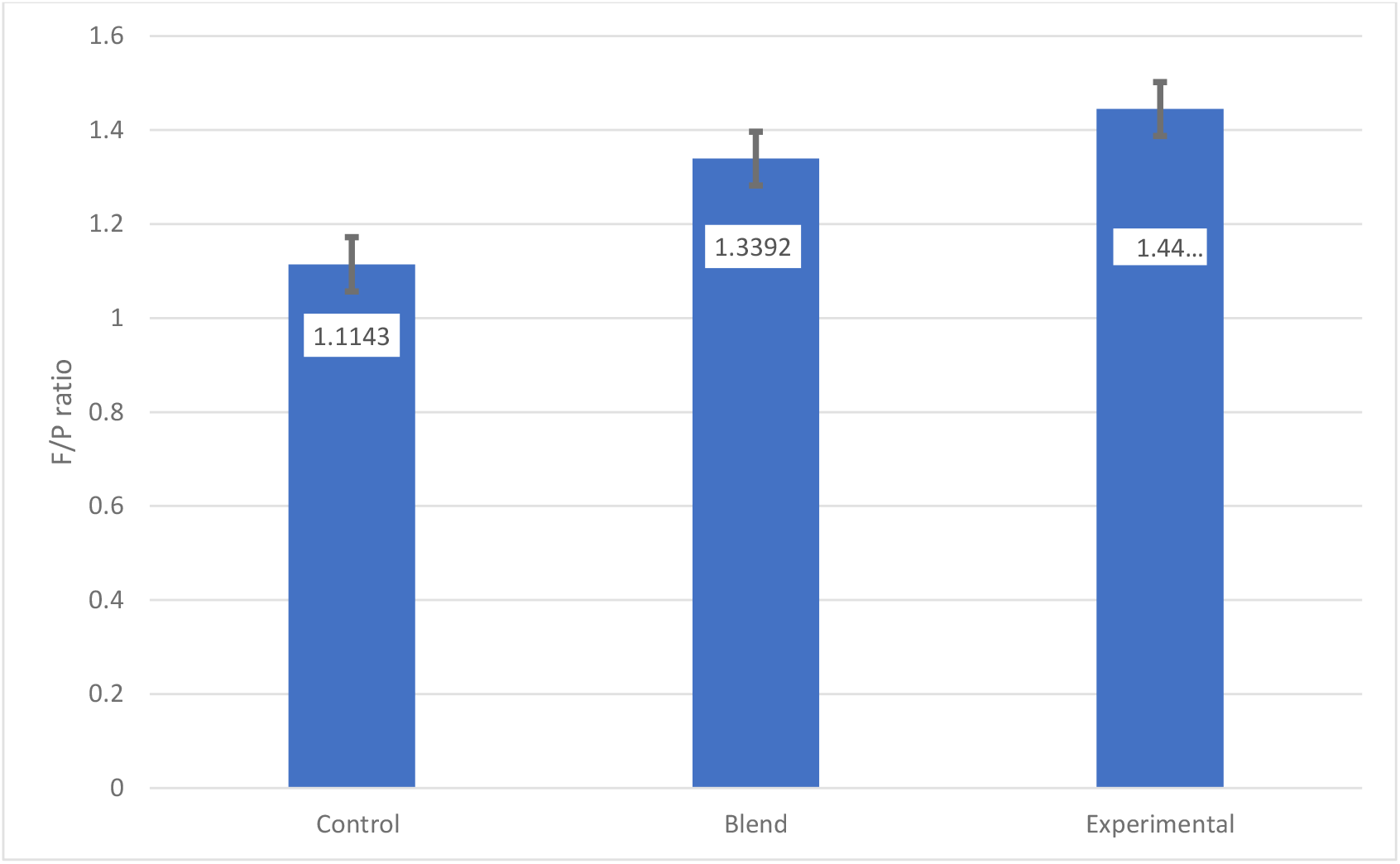
F/P concentration ratios with changing diets. Error Bars are +/- SE of each fitted mean.

The difference in the ratios confirms our hypothesis that the prebiotic action is primarily to enable the animal to abstract more **energy** from the diet (Jenkins 1993; Kristensen et al. 1998), but the increase in milk urea concentration, and the related increase in N ratio when the prebiotic was fed, suggests that the primary action of increasing degradation of fibre leads to increased availability of dietary N, which is then also used in some part for production of metabolic energy (Uddin et al. 2015).

### Washout analyses

The observed significant differences between treatments for milk composition may have been due to real diet effects, or due to non-random sampling. In human trials, it is often necessary to collect data for a period after treatment is completed to observe washout effects. Since the first evidence of statistical significance (for milk protein) was obtained two weeks after the trial commenced, it was presumed that a washout period of four weeks would be necessary, but sufficient, to permit milk composition to return to baseline. It is noted, however, that this period constitutes 10% of the average length of lactation in the trial herd, and that the period coincides with both physiological changes in the cow. In this trial, it also coincided with a period of warm, dry weather. Results were, therefore, considered in conjunction with changes in milk yield.

#### Milk yield

Milk yield differences persisted for a period after trial feeding ceased (Figure 5), then began to diminish. The data revealed a period of considerably increased variability from approximately one week after feeding ceased. Data for a further two weeks were collected.. Even after milk yield was corrected for baseline productivity, animals that had consumed the experimental diet maintained a continuing production advantage. (Control diet: 25.24L/day; Blend: 26.07L/day (103.3% of control); Experimental diet: 26.72L/day (105.9% of control) (p=0.000))

**Figure 5.**
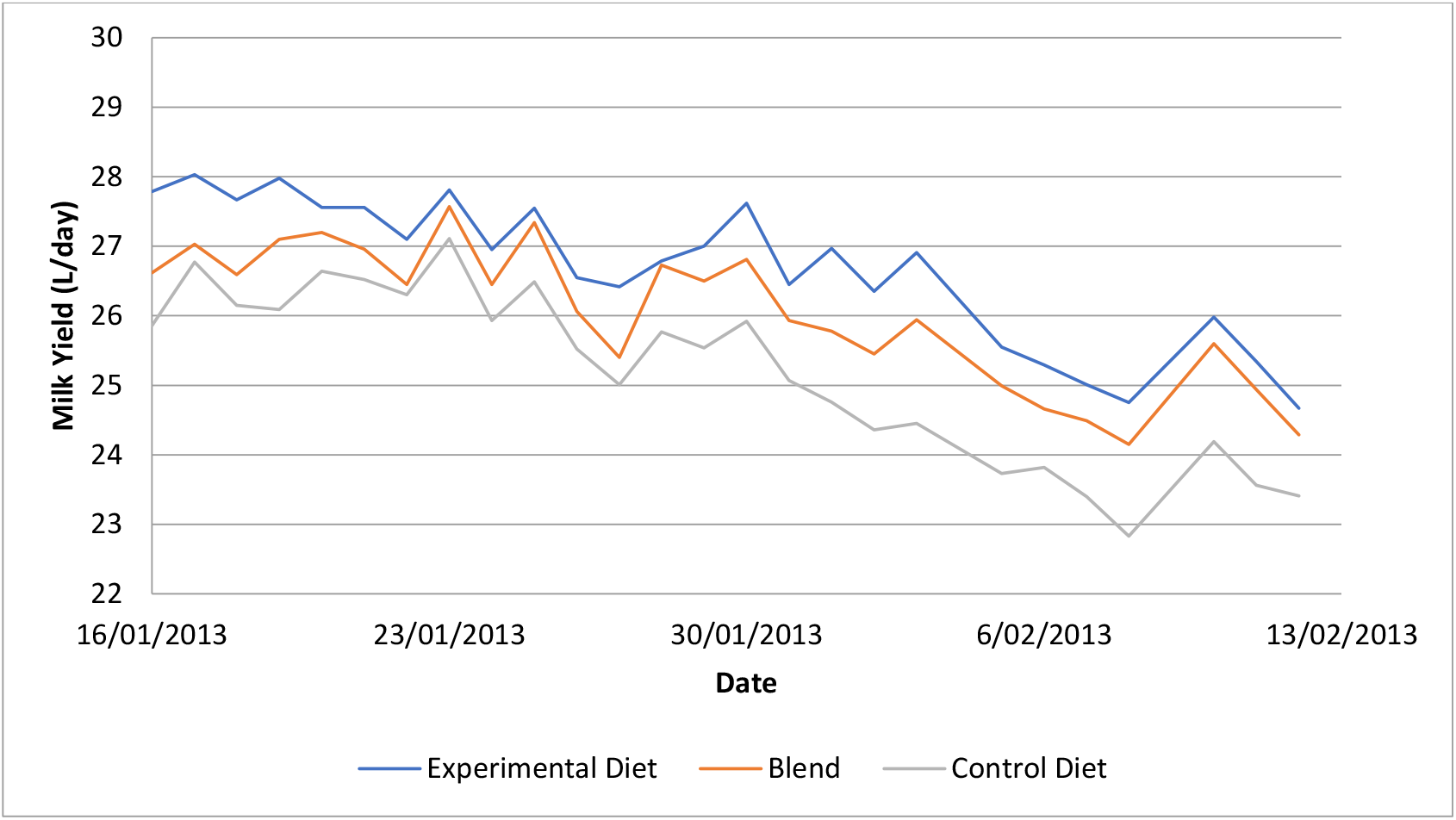
Daily milk production during washout. Prebiotic supplementation ceased on 16/01/2013 Raw data were corrected for variation in baseline milk production. Magnesium source for all animals in this phase was MgCl_2_ in drinking water.

#### Milk composition

With the exception of N ratio, significant differences between treatments for all characters measured had disappeared within one month of cessation of feeding (Table 5).

**Table 5.** Results of analysis of milk composition and solids yield.

| Character | Experimental diet<br>(% Control) | Blend<br>(% control) | Control diet | Significance (p) |
| --- | --- | --- | --- | --- |
| Fat (%) | 4.87 (125.2) | 4.91 (126.2) | 3.89 | 0.000 |
| Protein (%) | 3.33 (94.3) | 3.60 (102.0) | 3.53 | 0.008 |
| Lactose (%) | 4.87 (98.4) | 4.87 (98.4) | 4.95 | 0.108 |
| NZ Solids (%) | 8.26 (111.2) | 8.51 (114.5) | 7.43 | 0.000 |
| Urea (mM/L) | 28.71(110.0) | 26.53 (101.6) | 26.11 | 0.090 |
| Log SCC* | 4.804 | 4.719 | 5.186 | 0.001 |
| N ratio | 8.70 (118.5) | 7.40 (100.8) | 7.34 | 0.013 |
| Milk Energy | 15.95 (116.2) | 15.92 (116.0) | 13.73 | 0.000 |
| Milk Solids Yield<br>(kg/day) | 2.47 (122.2) | 2.00 (99.3%) | 2.02 | 0.002 |

\*Ratio to control is not appropriate when reporting transformed data

**Table 6.** Results of analysis of milk composition and solids yield.

| Character | Experimental diet<br>(% Control) | Blend<br>(% control) | Control diet | Significance (p) |
| --- | --- | --- | --- | --- |
| Fat (kg/day) | 1.413 (120.9) | 1.182 (101) | 1.169 | 0.009 |
| Protein (kg/day) | 1.005 (100) | 0.898 (89.4) | 1.005 | 0.001 |
| Urea (mM/L) | 28.71(110.0) | 26.53 (101.6) | 26.11 | 0.090 |
| Log SCC* | 4.804 | 4.719 | 5.186 | 0.001 |
| Milk Solids Yield<br>(kg/day) | 2.47 (122.2) | 2.00 (99.3%) | 2.02 | 0.002 |

\*Ratio to control is not appropriate when reporting transformed data

**Table 6.** Effect of treatment on milk composition during washout (samples collected fortnightly from 16/01/2013 until 13/02/2013)

| Character | Experimental diet<br>(% Control) | Blend<br>(% control) | Control diet | Significance (p) |
| --- | --- | --- | --- | --- |
| Fat (%) | 5.31 (128.5) | 5.25 (127.1) | 4.129 | 0.287 |
| Protein (%) | 3.50 (93.4) | 3.60 (103.4) | 3.75 | 0.094 |
| Lactose (%) | 4.78 (99.4) | 4.74 (98.6) | 4.82 | 0.711 |
| NZ Solids (%) | 8.81 (111.8) | 9.13 (115.9) | 7.88 | 0.341 |
| Urea (mM/L) | 36.17(122.3) | 30.17 (102.0) | 29.57 | 0.107 |
| Log SCC* | 4.74 | 4.86 | 5.22 | 0.071 |
| N ratio | 10.38 (131.2) | 7.90 (99.8) | 7.92 | 0.030 |
| Milk Energy | 16.73 (118.6) | 16.56 (117.4) | 14.10 | 0.288 |

\*Ratio to control is not appropriate when reporting transformed data

### Trend analysis

One-way ANOVA of milk solids yields (corrected for LogSCC) at each sampling date indicated that no significant difference between treatments could be observed, despite the overall finding of a highly significant advantage from feeding the trial product over a four-month period. An analysis in which data from successive sampling dates were added sequentially was conducted, and the p value for the treatment mean monitored until it fell below the threshold value of 0.05. In this investigation, this threshold was reached once the data from the first four sample collections were added to the analysis. Thus, in this study, it appears that the prebiotic effect took between 6 and 8 weeks to become unequivocal. Thereafter, statistical significance and milk solids yield advantage were retained or improved.

### Other findings

Across the entire herd, no changes in feed allocation were made during the course of either trial, clearly implying that increased productivity was not due simply to increased appetite. Consequently, if current assumptions about the relationship between feed intake and methane emissions are well-founded (Ramin and Huhtanen 2013), it seems that using the prebiotic will provide a reduction in the methane intensity of milk solids production.

Methane synthesis in the ruminant occurs in response to elevated proton concentration (reduced pH) in the ruminant, as a means of mitigating rumen acidosis. This, in turn, occurs as a result of vigorous short chain fatty acid (SCFA) production due to ruminal microbial fermentation. It is logical to assume that reducing the level of SCFAs by any means will reduce the pressure to produce methane (and other proton sinks). The apparent significant increase in triglyceride production, as shown by increased milk fat production, is clearly an alternative sink for ruminal SCFAs, and thus, potentially, may be accompanied by reduced methane emission.

## Conclusions

There was no evidence of any ill effects on animal health and welfare when a daily supplement of 22.0g of elemental Mg from MgO was replaced with 8.0g from crude magnesium lactate and ∼2.0g from flocs of soluble protein. As previously observed, a supplement containing crude magnesium lactate provided for a small but significant increase in daily milk yield, even when variation in baseline milk production was accounted for through covariate analysis. There appeared to be an almost linear dose response, as indicated by the blended diet.

However, there were several important, unexpected outcomes. Firstly, the experimental diet provided a statistically significant 22.2% increase in milk solids when analyte concentrations were corrected for milk volume, whereas the blended diet produced a slightly lower yield of total solids compared with the control. This effect was achieved **despite** an apparent **reduction** in milk protein from the experimental diet relative to the control. Unlike the first trial, the experimental diet in the second trial was associated with a relative increase in milk urea, which may explain the fate of protein which otherwise would have been available for secretion in milk. However, the milk urea concentrations observed in the study are not indicative of a major diversion of dietary N to energy production^2^.

It was observed that feeding the experimental product led to a reduction in somatic cell count of approximately 60%, almost entirely due to elimination of frank mastitis.

Note that the results above are achieved in comparison with a diet including an alternative supplement of wheat grain. When AME contribution effects of that supplement are accounted for, the estimated milk solids yield increase rises to 28.0%.

## Acknowledgements

The authors gratefully acknowledge the provision of commercial herd resources by Southern Centre Dairies for the conduct of the two trials reported, and the independent supervision of the trial work by Michael Sheppad and his colleagues at Vet4Farm, Invercargill.

The data and conclusions drawn were reviewed intensively by Dr Bruce Thorrold and his colleagues at DairyNZ, Hamilton, leading to a significantly improved manuscript, and the authors are most grateful for his contribution.

It is with considerable regret that we withdraw Paul Tocker from the list of authors of this paper, because of his sudden demise immediately after pronouncing himself satisfied with the manuscript.

## Footnotes

1 Note that the process also fixed bioavailable magnesium in flocculated protein, and this provided an extra 2g of magnesium in each daily dose.

2 Subsequent studies on-farm (Coles and Rowarth 2024) indicate that feeding Knewe®-Mg prior to calving elicits better “springing” (udder development) which has provided capacity to secrete about 7% more milk protein. Enhanced udder capacity may account for higher milk yield in the first study, where Knewe®-Mg was fed prior to calving, as this would presumably allow for the production and secretion of higher absolute amounts of lactose, which would come at the expense of milk fat production.

